# Differential Requirement of DICER1 Activity during Development of Mitral and Tricuspid Valves

**DOI:** 10.1101/2022.01.10.475694

**Authors:** Shun Yan, Yin Peng, Jin Lu, Saima Shakil, Yang Shi, David K. Crossman, Walter H Johnson, Shanrun Liu, Joy Lincoln, Qin Wang, Kai Jiao

**Author notes:** Correspondence to: Kai Jiao. These two authors contributed equally to this work (Shun Yan, Yin Peng).

## Abstract

Mitral and tricuspid valves are essential for unidirectional blood flow in the heart. They are derived from similar cell sources, and yet congenital dysplasia affecting both valves is clinically rare, suggesting the presence of differential regulatory mechanisms underlying their development. We specifically inactivated *Dicer1* in the endocardium during cardiogenesis, and unexpectedly found that *Dicer1*-deletion caused congenital mitral valve stenosis and regurgitation, while it had no impact on other valves. We showed that hyperplastic mitral valves were caused by abnormal condensation and extracellular matrix (ECM) remodeling. Our single-cell RNA Sequencing analysis revealed impaired maturation of mesenchymal cells and abnormal expression of ECM genes in mutant mitral valves. Furthermore, expression of a set of miRNAs that target ECM genes was significantly lower in tricuspid valves compared to mitral valves, consistent with the idea that the miRNAs are differentially required for mitral and tricuspid valve development. Our study thus reveals miRNA-mediated gene regulation as a novel molecular mechanism that differentially regulates mitral and tricuspid valve development, thereby enhancing our understanding of the non-association of inborn mitral and tricuspid dysplasia observed clinically.

**Summary statement:** Although mitral and tricuspid heart valves are both derived from same cell sources in the atrioventricular region through similar initiation and remodeling processes, congenital dysplasia simultaneously affecting both valves is clinically uncommon. We show that DICER1 activity is only essential for mitral valve development whereas it is dispensable for other valves. Our study thus reveals miRNA-mediated gene regulation as a novel molecular mechanism that is differentially required to regulate mitral and tricuspid valve development, thereby enhancing our understanding of the non-association of inborn mitral and tricuspid dysplasia observed clinically.

## Introduction

Congenital heart diseases (CHDs) occur in as many as 1-5% of newborns and remain the leading noninfectious cause of morbidity and mortality for infants in developed countries (Clark et al., 2006; Hoffman, 1995; Hoffman and Kaplan, 2002; Onuzo, 2006). Malformations of valves represent major forms of CHDs, accounting for up to 30% of CHDs (Armstrong and Bischoff, 2004; Combs and Yutzey, 2009; LaHaye et al., 2014). Malformation of valves during cardiogenesis can also predispose valves to late onset valvular diseases in adults (LaHaye et al., 2014; O’Donnell and Yutzey, 2020). Therefore, understanding the molecular, cellular and genetic mechanisms underlying normal valvulogenesis and their contributions to inborn valve diseases has a strong translational significance.

Cardiac valves are essential structures to ensure unidirectional blood flow in the heart. Valvulogenesis in mouse embryos is initiated with regional expansion of extracellular matrix (ECM) in the atrioventricular (AV) canal region and the outflow tract (OFT) region between E9.0 to E10.0 (Armstrong and Bischoff, 2004; Butcher and Markwald, 2007; Combs and Yutzey, 2009; de Vlaming et al., 2012; LaHaye et al., 2014; Lin et al., 2012; O’Donnell and Yutzey, 2020). Shortly after the accumulation of cardiac jelly, a group of endocardial cells in the AV cushion and OFT conal cushion (the proximal region of the OFT) respond to inductive signals released from the overlying myocardium and undergo epithelial-mesenchyme-transition (EMT) to invade into the ECM of cushions (*ibid*.). In the AV cushions, mesenchymal cells are derived from endocardial cells and therefore deletion of a gene in endocardial cells prior to EMT will lead to permanent deletion of the gene in cushion mesenchymal cells. Cellularized cushions are subjected to sophisticated and well-controlled remodeling processes, including condensation (increase in cell density), elongation and ECM remodeling, and eventually become mature thin valve leaflets (*ibid*). The endocardial-derived mesenchymal cells differentiate into interstitial cells residing in the ECM of valves and are essential for valve homeostasis.

The mitral and tricuspid valves separate atria and ventricles of the left and right sides of the heart, respectively. While they both are derived from similar cell sources through similar initiation and maturation processes, it is highly uncommon in human CHD patients that both AV valves are affected simultaneously. In a previous study that examined co-occurrence of different forms of CHDs in >3000 individuals with clinically confirmed phenotypes, it was found that the Odds Ratio (OR) between abnormal tricuspid valves and abnormal mitral valves was equal to 1, indicating that abnormalities of the two AV valves occur independently of one another in patients (Ellesoe et al., 2018). It is also well known that congenital polyvalvular heart disease, which affects more than one valve simultaneously, mainly occurs in patients with chromosomal abnormalities, such as trisomy 13 and trisomy 18, and is rarely observed in people with normal chromosomes (Bartram et al., 2001). These clinical observations cannot be easily explained by our current knowledge regarding the general mechanisms regulating AV valvulogenesis, as abnormalities in these mechanisms are expected to affect formation of both AV valves. Rather, the non-association between mitral and tricuspid valve inborn dysplasia strongly suggests the presence of regulatory mechanisms that differentially regulate development of the two AV valves. These mechanisms are vulnerable to detrimental genetic/environmental factors and defects affecting the differential regulatory mechanisms, rather than the well-studied general mechanisms, and are likely the major cause of AV valvular anomalies observed in human CHD patients. Our current knowledge regarding the molecular mechanisms differentially regulating mitral and tricuspid valve development in embryonic hearts remains largely limited.

Micro RNAs (miRNAs) are non-coding small RNA molecules that regulate numerous biological processes including cardiovascular development (Chen and Wang, 2012; Cordes and Srivastava, 2009; Malizia and Wang, 2011; O’Brien et al., 2018; Yan and Jiao, 2016). MiRNA genes are first transcribed by RNA polymerase II/III to generate Pri-miRNAs, which are then processed by DROSHA in the nucleus before being transported into the cytoplasm. In the cytoplasm, Pri-miRNAs are catalyzed by the RNase III ribonuclease DICER to generate the mature miRNAs, which are single stranded RNA composed of 22-25 nucleotides. Mature miRNAs act on complementary 3’ untranslated regions (3’ UTRs) of target mRNAs to downregulate their stability and translation efficiency (*ibid*.). DICER is thus required for the production of most functional miRNAs and deletion of *Dicer1* blocks miRNA biosynthesis. The essential role of miRNA mediated gene regulation on heart development has been well demonstrated by conditional inactivation of *Dicer1* in cardiomyocytes (Chen et al., 2008; Peng et al., 2014; Saxena and Tabin, 2010; Zhao et al., 2007), neural crest cells (Huang et al., 2010a; Huang et al., 2010b; Nie et al., 2011; Zehir et al., 2010), and epicardial cells (Singh et al., 2011). However, the function of miRNAs in the endocardium and valves during mammalian cardiogenesis have been largely overlooked in the literature.

In this study, we attempted to test how blocking miRNA biosynthesis by deletion of *Dicer1* in the endocardium affects valvulogenesis using *Nfatc1-Cre*. Endocardial deletion of *Dicer1* caused lethality within 3 days after birth, indicating that endocardial expression of *Dicer1* is essential for the survival of newborn mice. Furthermore, we unexpectedly found that deletion of *Dicer1* led to mitral valve stenosis and regurgitation in perinatal hearts, while it had a minor impact on other valves including tricuspid valves in the same mice. Our study therefore suggests, for the first time, that miRNA mediated gene regulation is differentially required for mitral and tricuspid valve development. These results will help us understand the molecular basis for the clinical observations that congenital dysplasia rarely affects both AV valves simultaneously and further suggest that defects in miRNA mediated gene regulation can be a potential etiological factor for inborn mitral valve stenosis and regurgitation.

## Results and Discussion

### 1. Endocardial deletion of *Dicer1* led to mitral valve stenosis and regurgitation

To understand how blocking miRNA biosynthesis affects cardiac valvulogenesis, we inactivated *Dicer1* in endocardial cells using *Nfatc1-Cre* mice, which can efficiently inactivate target genes in endocardial cells before EMT is initiated (Peng et al., 2016; Wu et al., 2012). We crossed *Nfatc1-Cre/Dicer1*^*loxp/+*^ male mice with *Dicer1*^*loxp/loxp*^ female mice (Cobb et al., 2005; Peng et al., 2014) to obtain conditional knockout (cKO, *Nfatc1-Cre/Dicer1*^*loxp/loxp*^) and control (*Dicer1*^*loxp/+*^ or *Dicer1*^*loxp/loxp*^) animals at various developmental stages. Our quantitative reverse transcription polymerase chain reaction (qRT-PCR) results confirmed that expression of *Dicer1* transcripts was efficiently inactivated in both mitral and tricuspid valve primordium at E11.5 (Sup Fig. 1). Mutant mice were recovered at the Mendelian ratio (∼25%) until postnatal day 0 (P0). Starting from P1, the percentage of live mutant mice was significantly reduced and no mutants survived to P3 (Fig. 1A), indicating that endocardial deletion of *Dicer1* is incompatible with postnatal survival. Whole-mount examination of P1 hearts showed that the left atrium (LA) of mutant hearts was dilated and was heavily congested with blood (Fig. 1B). Section examination showed that the leaflets of mitral valves in mutant hearts were hyperplastic and abnormally shaped, while the morphology of other valves (including tricuspid and semilunar valves) were not overtly different from control mice (Fig. 1C-E). The orifice of mutant mitral valves was reduced and blood cells were trapped in the LA. To examine further the functional relevance of the abnormalities in mutant mitral valves, we examined mitral flow in P1 mice by echocardiography color Doppler (Fig. 1F, G). The mitral flow rate was significantly increased in mutant samples, consistent with the observed hyperplastic phenotype and reduced orifice of mitral valves. Furthermore, regurgitation was observed in all mutant samples examined. We thus concluded that deletion of *Dicer1* in endocardial cells caused congenital mitral valve stenosis and regurgitation while having a minor impact on other valves. This is a somewhat unexpected result, as mitral and tricuspid valves are formed through similar processes and it was expected that blocking miRNA biosynthesis would affect development of both valves. Our reporter and qRT-PCR analyses (Sup. Fig. 1) excluded the possibility that *Dicer1* was only efficiently inactivated in mitral vales but not in tricuspid valves. Therefore, our data clearly demonstrated the differential requirement of *Dicer1* between mitral and tricuspid valves during their development.

**Figure 1.**
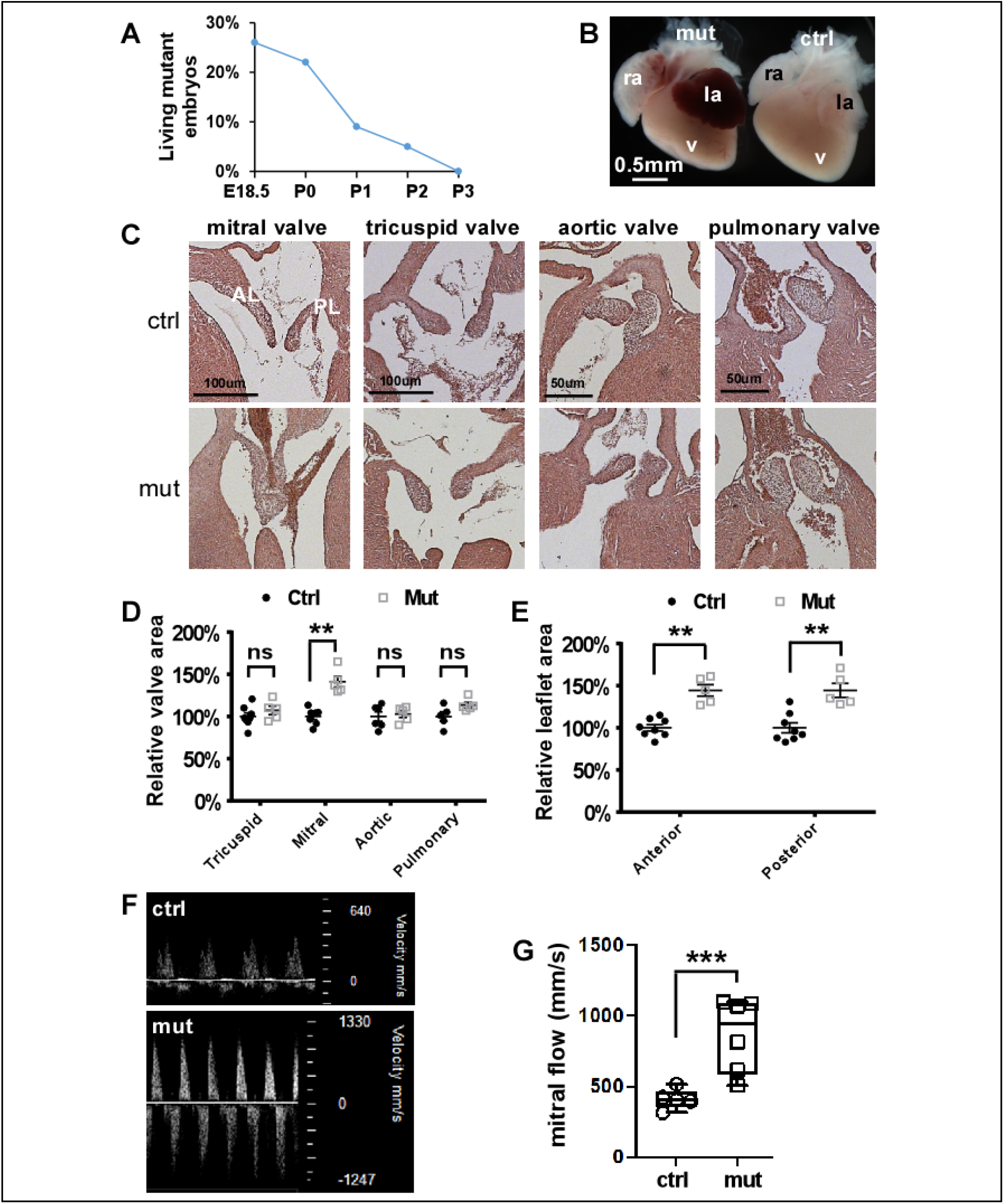
Endocardial-deletion of *Dicer1* led to mitral valve stenosis and regurgitation. **(A)** The percentage of living mutant embryos (*Nfac1-Cre*;*Dicer1*^*loxp/loxp*^) from the cross between male *Nfac1-Cre*;*Dicer1*^*loxp/+*^ and female *Dicer1*^*loxp/loxp*^ mice. Data were calculated from 10 litters of mice. **(B)** Whole mount examination of P1 hearts. The left atrium of the mutant heart was dilated with blood trapped inside. **(C)** Morphological examination of valves at P1. The arrow shows the reduced size of the orifice of the mutant mitral valve. **(D)** Quantification of leaflet area from different valves. P1 hearts were sectioned to 10um thickness and stained with HE. The area of leaflets in each section were measured using ImageJ, and total areas were added together. The area of control leaflets was set at 100%. Data were averaged from 5 mutant and 6 control hearts with error bars showing standard error (SEM). **(E)** Quantification of anterior and posterior leaflets of mitral valves from control (n=8) and mutant hearts (n=5). **(F, G)** Mitral valve flow measurement through echocardiography color Doppler. Mitral flow velocity was measured by localizing the sample volume to basal mitral valve inflow and optimizing position. In panel F, representative images show that maximal mitral valve inflow velocity was higher in mutant hearts compared to control hearts. The white arrow indicates regurgitation. Panel G shows summary data of mitral valve flow velocity from WT (n=5) and mutant (n=6) hearts. In panel A, **: *p<0*.*01*, Chi-square test. In other panels, **: *p<0*.*01*, ***: *p<0*.*001*, unpaired, two tailed Student’s t test; ns: not significant. la: left atrium, ra: right atrium, v: ventricle.

### 2. Deletion of *Dicer1* caused mitral valve remodeling defects in perinatal hearts

To determine at which stage mutant mitral valves started to become hyperplastic, we examined valve development starting from E9.5. We did not observe valve abnormalities until E17.5 and hyperplastic mitral valves became overt in mutant hearts at E18.5 (Fig. 2A, B). We next attempted to reveal the cellular mechanism underlying hyperplastic mitral valves by examining cell proliferation, apoptosis and remodeling. We did not observe significantly altered cell proliferation and death in mutant mitral valves at E16.5-18.5 (Fig. 2C, D, Sup. Fig. 2). Therefore, deletion of *Dicer1* did not affect mitral valve cell proliferation or survival. We next examined the cell density in mitral valves, and revealed that the cell density (number of cells normalized against the area) in the middle and distal regions of mitral valves was significantly decreased in mutants (Fig. 2E), suggesting that deletion of *Dicer1* impaired cell condensation of mitral valves. Our further *in vitro* collagen gel contraction assay showed that the cells isolated from mutant mitral valves were less capable of contracting collagen gels (Fig. 2G), suggesting that their capacity to remodel extracellular matrix (ECM) was reduced. Our data collectively suggest that *Dicer1* is required for normal remodeling of mitral valves at the perinatal stage and the reduced condensation may be the major factor causing hyperplastic mitral valves in mutants.

**Figure 2.**
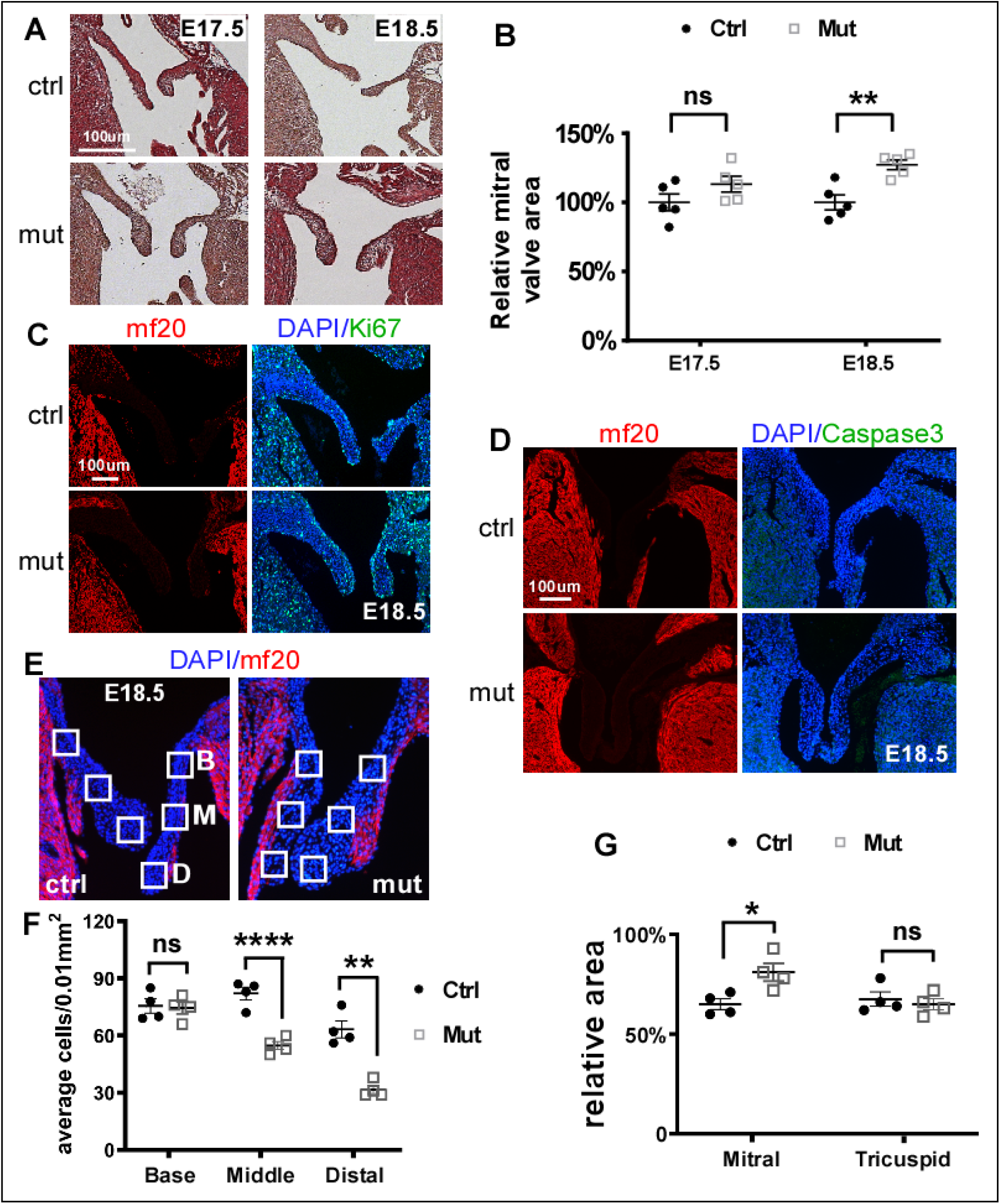
Endocardial-deletion of *Dicer1* led to defects in condensation and ECM remodeling in mitral valves. **(A)** Mitral valves of control (ctrl) and mutant (mut) hearts at E17.5 and E18.5. **(B)** Quantification of areas of mitral valves at E17.5 and E18.5 (n=5). **(C)** Sections of control and mutant hearts at E18.5 covering the mitral valve area were stained with antibodies against cardiac myosin heavy chain (mf20, red) and Ki67 (green). Total nuclei were visualized with DAPI staining (blue). **(D)** Sections of control and mutant hearts at E18.5 covering the mitral valve area were stained with antibodies against cardiac myosin heavy chain (mf20, red) and cleaved CASP3 (green). **(E, F)** Cardiac sections covering the mitral valve area were stained with an mf20 antibody (red) and DAPI (DAPI) in panel E. The boxed areas indicate the base (B), middle (M) and distal (D) positions of mitral valves that were used to measure cell density. At least 5 sections of each heart from 4 hearts were examined and measured. The data are shown as mean ± SEM in panel F. We did not observe a difference between anterior and posterior leaflets, and thus data from both leaflets were combined. **(G)** Collagen gel contraction assay using cells isolated from mitral valves of control and mutant hearts at E18.5. Data were averaged from 4 independent cultures with error bars indicating SEM. 48-hours post incubation, the areas of gels without cells were set at 100%. *: *p<0*.*05;* **: *p<0*.*01*, ****: *p<0*.*0001*, unpaired, two tailed Student’s t test; ns: not significant.

### 3. Expression of multiple extracellular matrix genes is impaired in mutant mitral valves

To understand the molecular defects in mitral valves caused by *Dicer1*-deletion, we performed drop-let based single cell RNA-Sequencing (scRNA-Seq) using the mitral valves dissected from control and cKO mice at E17.5, a stage when the morphological defect was not overt. Performing studies using cells from stage E17.5 increased the likelihood that we would reveal the molecular alterations that cause the hyperplastic valve phenotype, rather than those that result from the defect. We were able to acquire 10,434 and 6014 cells from control and mutant samples, respectively. Using the Uniform Manifold Approximation and Projection (UMAP) dimension reduction technique (Becht et al., 2018), these cells could be grouped into at least 29 clusters (Sup. Fig. 3). Based on known molecular markers of cells in mouse developing hearts (Hill et al., 2019; Li et al., 2016), we were able to reveal various cardiac cell types including cardiomyocytes, endocardial cells, endothelial cells, epicardial cells, and valve mesenchymal cells (Fig. 3A, B, Sup. Fig. 4). In addition, we also identified red blood cells, white blood cells and lymphatic endothelial cells. We then focused on mesenchymal cells, which are largely derived from AV cushion endocardial cells and are the major cell type in mitral valves undergoing remodeling. We compared gene expression between control and mutant mesenchymal cells, and revealed that 169 genes whose expression was significantly altered by at least 25% with adjusted *p<0*.*01*. We next performed gene ontology (GO) term enrichment analysis using Metascape (Zhou et al., 2019), and revealed that “Extracellular matrix organization” and “Extracellular structure organization” are the two most significantly affected pathways (Fig. 3C, Sup. Table 1). These pathways contain >20 genes that encode the ECM proteins and examples of 10 genes are shown in the volcano chart in Fig. 3D. Therefore, results from our molecular examination suggest abnormal expression of ECM genes as a major molecular defect contributing to the remodeling defect in mitral valves.

ECM plays critical roles in regulating valve formation and homeostasis (Kodigepalli et al., 2020; O’Donnell and Yutzey, 2020). Mutations in multiple ECM genes, such as *COL1A1, COL3A1, FLNA*, and *TNXB*, can lead to valve diseases in human patients (Brady et al., 2017). In a recent study, a mutation in a ciliogenesis regulatory gene, *DZIP1*, was found to cause mitral valve prolapse and it was further determined that primary cilia acts through regulating ECM distribution and organization to promote normal mitral formation and homeostasis (Toomer et al., 2019). We now show with clear *in vivo* mouse genetic evidence that *Dicer1* is required for expression of many ECM genes. The proteins encoded by these genes cover many components of the ECM in valves, including collagen, core protein of proteoglycans, elastic fibers, fibronectin and laminins, suggesting the broad activity of miRNAs in regulating ECM genes in mitral valves.

The hemodynamics of the left and right sides of postnatal hearts are significantly different from each other (Arvidsson et al., 2017), and mitral and tricuspid valves adapt to different hemodynamic forces to function. ECM proteins play a major role in determining the biomechanics of heart valves (Kodigepalli et al., 2020). The hemodynamic difference between left and right sides of embryonic hearts and its potential role in regulating valve development have not been well-defined. It will be interesting in future studies to test if miRNA mediated regulation of ECM gene expression is involved in mediating the hemodynamic forces to shape mitral valve development in embryonic hearts.

In the next step, we performed trajectory and pseudotime analysis on valve mesenchymal cells using Monocle 3.0. The mesenchymal cells (control + mutant) could be further grouped into 5 subclusters (Sup. Fig. 5). Based on pseudotime ordering, the trajectory started with subcluster 1, which sequentially gave rise to cells in subclusters 2-5 (Fig. 3E). Cells in subcluster 1 are least matured, and consistently, in the UMAP chart, they are closely adjacent to endocardial cells (Fig. 3A), which are the precursors of AV valve mesenchymal cells. An increased percentage of cells in subcluster 1 was observed in the mutant sample compared to wild type (14% vs. 5%) (Fig. 3E), suggesting that mutant mitral valves contain more immature mesenchymal cells. Interestingly, when performing trajectory analysis separately on control and mutant cells, we noticed that the trajectory between subcluster 1 and other subclusters was broken in control samples (Sup. Fig. 6). This result suggests that in wild type mitral valves at E17.5, differentiation of cells in subcluster 1 into cells in other subclusters already stopped, whereas such a trajectory still existed in mutant valves. Data from trajectory and pseudotime analyses suggest that *Dicer1* is required for proper maturation of mitral valves at the perinatal stage.

**Figure 3.**
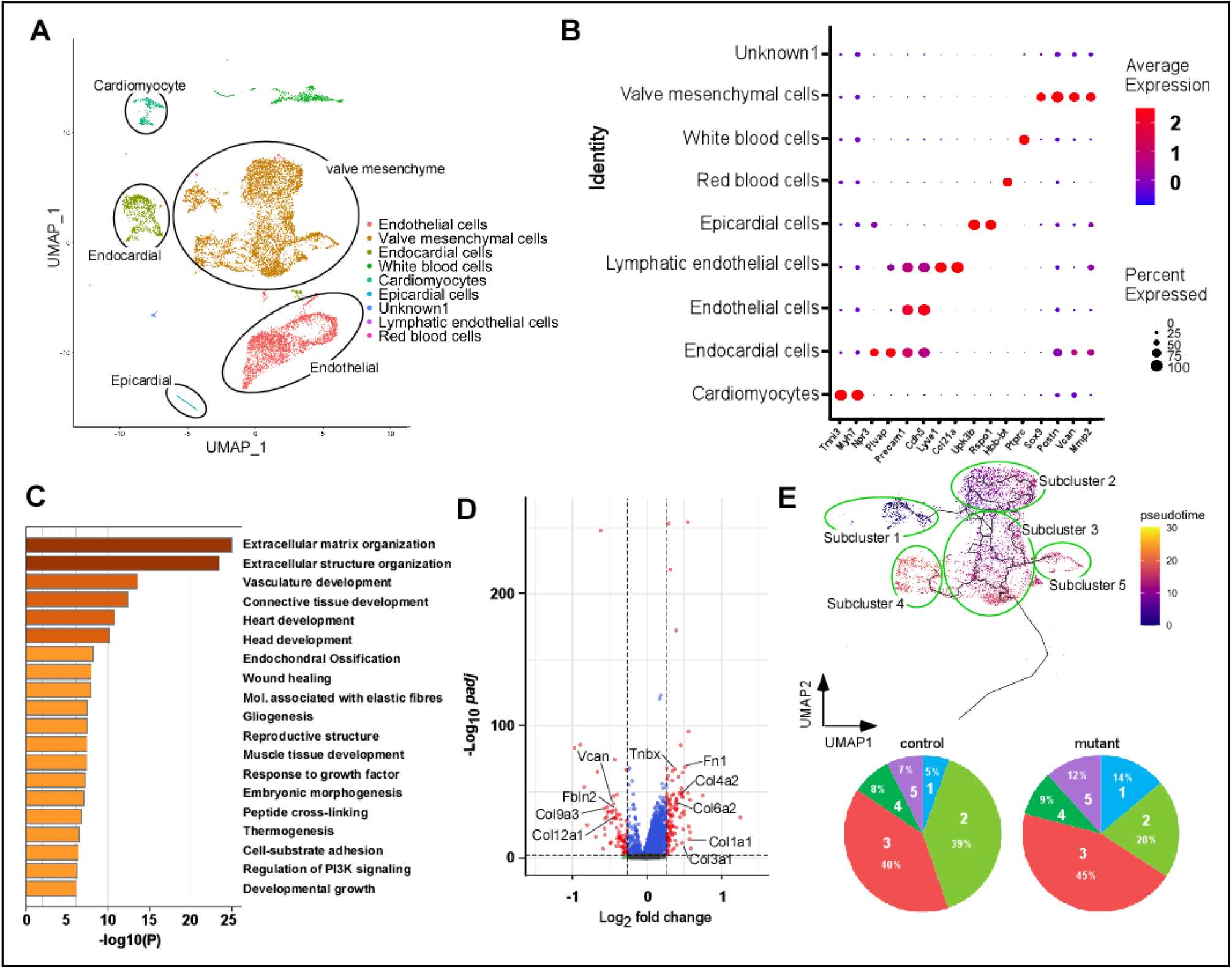
ScRNA-Seq analysis of mitral valves of E17.5 hearts. **(A)** The mitral valves from control and mutant hearts at E17.5 were isolated to obtain single cell suspensions, which were then subjected to scRNA-Seq analysis. This chart shows UMAP projection of various cell types. Clusters corresponding to cardiomyocytes, epicardial cells, endocardial cells, endothelial cells and valve mesenchymal cells are indicated with circles in the chart. **(B)** The Dotplot chart shows expression of representative marker genes of different cell types in hearts across all clusters. **(C)** The differentially expressed genes between control and mutant valves were subjected to GO term enrichment analysis using Metascape. The terms involved in ECM organization and extracellular structure organization were ranked as top 1 and 2, respectively. **(D)** The volcano chart shows differentially expressed genes with the X-axis showing the fold of alteration and the Y-axis showing the Benjamini-Hochberg adjusted P value. The dotted lines show the threshold (adjusted *p<0*.*05*, expression altered by at least 25%). 10 ECM genes that are differentially expressed are indicated. **(E)** Trajectory and pseudotime analysis of valve mesenchymal cells. In the top panel, the mesenchymal cells were further divided into 5 subclusters using Monocle 3.0. The pseudotime is indicated with the color and the trajectory pathways are shown with black lines in the UMAP chart. The pie charts in the bottom panel show the percentage of cells in each subcluster in control and mutant samples. The percentage of cells in subcluster 1, which are least matured cells, is increased from 5% in control samples to 14% in mutant samples.

### 4. Expression of *Col1a1, Col3a1* and *Tnxb* in mitral valves is directly regulated by miRNAs

Among the ECM genes that were upregulated by *Dicer1*-deletion, we decided to focus on *Col1a1, Col3a1* and *Tnxb*. Mutations in these three genes are associated with defects in mitral valves (Brady et al., 2017). We first verified increased expression of these genes at the protein level in mutant mitral valves using immunostaining analysis (Fig. 4A). We then performed sequence analysis using miRNA target prediction programs including TargetScan and miR Walk, and identified multiple potential miRNA targets in the 3’ UTRs of the three genes (Sup. Fig. 7). To test if the 3’ UTRs of the three genes are targeted by relevant miRNAs, we inserted the 3’ UTRs of these genes into a reporter vector and performed reporter analysis using tsA58-AVM cells, which are temperature sensitive immortal cells derived from AV cushion mesenchymal cells (Peng et al., 2016). As shown in Fig. 4B, mimics of miR98-5p and miR29a reduced the activity of reporters derived from the 3’UTRs of *Col1a1* and *Col3a1* in a dose dependent manner and the mimic of miR30c significantly reduced the activity of the *Tnxb* reporter. Our results thus suggest that transcripts of *Col1a1* and *Col3a1* can be regulated by miR98/Let7 and miR29 in endocardium-derived valve mesenchymal cells and the transcript of *Tnxb* can be targeted by miR30. On the other hand, mimics of miR148a and miR196b had no effect on *Col3a1* and *Tnxb* reporters, arguing against their direct role in regulating transcripts of *Col3a1* and *Tnxb* (Sup. Fig. 8). To further test if mitral valves express corresponding miRNAs that target transcripts of *Col1a1, Col3a1* and *Tnxb*, we performed qRT-PCR analysis and confirmed that AV valve mesenchymal cells express multiple family members of miR98/let7, miR29 and miR30, (Fig. 4C). Of particular interest, we noticed that the expression level of multiple miRNAs of these families was significantly lower in tricuspid valves than in mitral valves. The reduced expression of miRNAs in tricuspid valves suggests that miRNA-mediated gene regulation plays a less significant role in regulating ECM genes compared to mitral valves. This observation helps explain the absence of overt defects in cKO tricuspid valves. We speculate that in mutant tricuspid valves, expression of a set of miRNAs that target ECM genes is lower than in mitral valves and the loss of DICER1 can be sufficiently compensated by other regulatory mechanisms to maintain proper ECM gene expression. On the contrary, in mitral valves, miRNA-mediated regulation of ECM gene expression plays a more substantial role than that in tricuspid valves, and compensatory mechanisms cannot sufficiently restore the normal level of ECM proteins in mutant mitral valves, leading to mitral valve defects. Based on our mouse data, we further predict that mutations in miRNA genes or genes involved in miRNA biosynthesis preferentially affect mitral valve development and act as potential etiological factors for inborn mitral valves dysplasia. Therefore, miRNA-mediated gene regulation may serve as a specific therapeutic target for congenital mitral valve defects.

**Figure 4.**
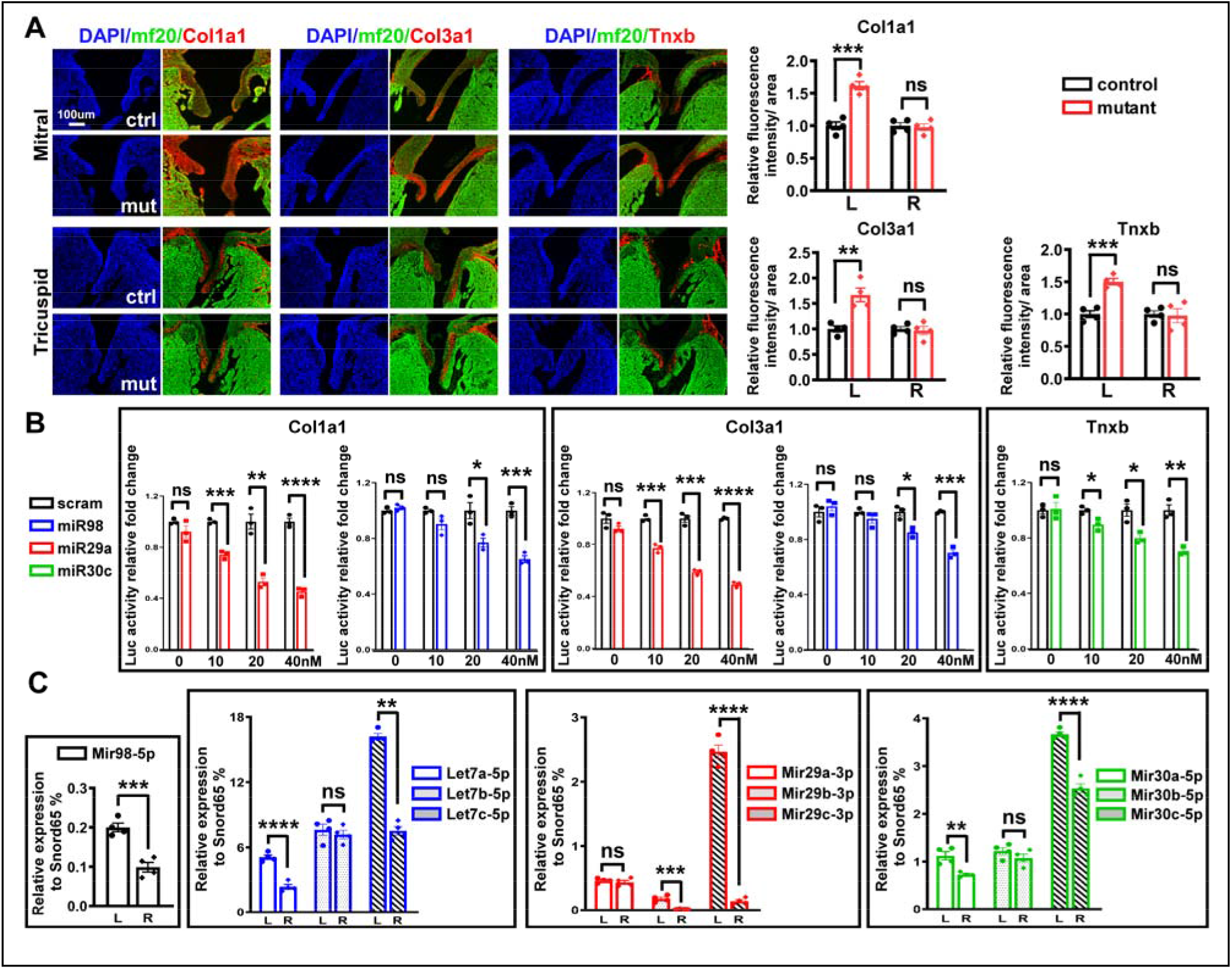
*Col1a1, Col3a1* and *Tnxb* are targets of a group of miRNAs in developing mitral valves. **(A)** Cardiac sections (E18.5) were stained with antibodies as indicated. Total nuclei were visualized with DAPI staining (blue). Areas covering mitral and tricuspid valves are shown. The charts in the right panels show the quantification of immunofluorescent intensity normalized against area measured using ImageJ. Data were acquired from 4 independent hearts of each genotype and for each heart at least 3 different sections were measured. The intensity of control samples are set to 1.0. Data are shown as mean ± SEM. **(B)** Luciferase reporter analysis was performed using various reporter constructs harboring the 3’ UTA region of indicated genes in tsA58-AVM cushion mesenchymal cells. Cells were co-transfected with reporter constructs and various miRNA mimics at different doses, and the value of cells transfected with scrambled miRNA was set at 1.0. **(C)** RNA samples (with enrichment of small sizes) were purified from cells isolated from wild type tricuspid and mitral valves. QRT-PCR was then performed to examine expression of various miRNAs. The expression level of miRNAs was normalized against Snord65. L: left side, mitral valve; R: right side, tricuspid valve. *: *p<0*.*05;* **: *p<0*.*01*; ***: *p<0*.*001*; ****: *p<0*.*0001*, unpaired, two tailed Student’s t test; ns: not significant.

In summary, we unexpectedly observed that *Dicer1* is differentially required for the development of mitral and tricuspid valves, which are both derived from AV cushions through similar initiation and remodeling processes. *Dicer1* is essential for proper expression of multiple ECM genes in mitral valves, and endocardial deletion of *Dicer1* impaired maturation and remodeling of mitral valves. Our results reveal miRNA-mediated gene regulation as a novel regulatory mechanism that is differentially required for mitral and tricuspid valve development, and help explain the clinical observation that co-occurrence of congenital defects affecting both AV valves is uncommon. These data will ultimately contribute to development of novel clinical applications toward diagnosis and treatment of AV valve defects.

## Materials and Methods

### 1. Mouse strains and maintenance

This study was carried out in accordance with the Guide for the Care and Use of Laboratory Animals published by the US National Institutes of Health (NIH Publication no. 85–23, revised 2011). All protocols were approved by the Institutional Animal Care and Use Committee (IACUC) at the University of Alabama at Birmingham. Mice were euthanized by inhalation of CO2 followed by cervical dislocation. The *Nfatc1-Cre* and *Dicer1*^*loxp/loxp*^ mouse lines were described previously (Cobb et al., 2005; Peng et al., 2014). Mice were maintained on the C57BL/6 background. *Nfatc1-Cre;Dicer1*^*loxp/+*^ male mice were crossed with *Dicer1*^*loxp/loxp*^ females to obtain control and cKO mice at different developmental stages. Staged littermates of embryos were collected, with stage E0.5 defined by the presence of a copulation plug. Primers and PCR conditions for genotyping were described previously (Peng et al., 2014; Yan et al., 2020).

### 2. Histological and immunostaining analyses

All experiments were performed as described in previous studies (Liu et al., 2014; Peng et al., 2016; Yan et al., 2020). For hematoxylin and eosin (HE) staining, embryos and perinatal hearts were fixed in 4% paraformaldehyde overnight at 4°C. The next day, tissues were washed in PBS, dehydrated, and embedded in paraffin. 10um paraffin sections were stained with HE for morphological examination. For Immunofluorescence, paraffin sections were dewaxed, rehydrated, and treated with citrate buffer for antigen retrieval. After being blocked with 10% serum in TBST at room temperature for 1-3 hours, slides were incubated with a primary antibody overnight at 4°C, and then incubated with an Alexa-conjugated secondary antibody (ThermoFisher) at room temperature for 1 hour. Finally, slides were briefly stained with 4′,6-diamidino-2-phenylindole (DAPI, ThermoFisher) to reveal the nuclei. When needed, the intensity of the immunostaining signal was amplified by the TSA Plus Cyanine 3.5 System (PerkinElmer). Immunostained samples were observed using a Zeiss Axio fluorescent microscope. The primary antibodies used for immunostaining were purchased from Iowa Hybridoma Bank (MF20, 1/500), Cell Signaling (cleaved Caspase3, #9661, 1/500), Abcam (Ki67, #ab15580, 1/500; COL1A1, #ab34710, 1/100), Novus Biologicals (COL3A1, #NB600-594, 1/100), and R&D systems (TNXB, #AF6999, 1/100).

### 3. Measurement of the area and cell density of valve leaflets

Embryonic hearts at various stages were frontally sectioned to a 10um thickness. The area of each leaflet of all sections was measured using ImageJ, and then added to calculate the total area for each valve leaflet. The sections were stained with an antibody against cardiac myosin heavy chain (MF20) to avoid inclusion of cardiomyocytes in the measurement. Data were averaged from at least 5 independent hearts of each genotype and the total area for controls was set at 100%. To measure cell density in mitral valve leaflets, cardiac sections (E18.5) were stained with an MF20 antibody (against cardiac myosin heavy chain) to view cardiomyocytes and with DAPI to view total nuclei. Staining with the MF20 antibody helped us to exclude cardiomyocytes in our counting. Cell density of the base, middle and distal regions of anterior and posterior leaflets of mitral valves were calculated as total number of nuclei normalized against the area examined. At least 5 sections of each heart from 4 hearts of each genotype were quantified. No difference between anterior and posterior leaflets was observed and thus data from both leaflets of each mitral valve were combined.

### 4. Measurement of mitral valve flow by echocardiography

Mitral flow was identified in control and mutant neonatal mice (P1) by echocardiography color Doppler and velocity was measured by pulsed wave Doppler using the Vevo 3100 ultra-high-resolution ultrasound system (Fuji, Visual Sonics, Canada) with MX400 linear array mouse transducer. Pups were lightly anesthetized under inhaled isoflurane anesthesia (1.5-2.0% isoflurane in 100% O2) and maintained on a heated platform (37°C) during echocardiographic analysis. Mitral valve inflow was localized in parasternal long axis 2 chamber view in the basal aspect posterior to the aortic valve and velocity measured after optimizing sample volume placement by pulsed wave Doppler. Maximal mitral flow velocity was determined in each pup and used in group analysis.

### 5. Collagen gel contraction, Luciferase reporter and qRT-PCR analyses

Collagen analysis was performed using the CytoSelect Cell Contraction Assay Kit (Cell Biolabs, Inc). Mitral valves were dissected from control and mutant hearts at E18.5 and were digested with 1mg/ml collagenase and dispase (Sigma-Aldrich, #10269638001) at 37°C for 4 minutes followed by 0.25% trypsin at 37°C for 10 minutes to acquire single cell suspensions. Cells from 3-4 mitral valves of the same genotype were pooled to acquire 10k cells for one well of a 96-well plate. Cells were mixed with the collagen solution following instructions provided by the vendor. Plates were then incubated in a cell culture incubator (37°C with 5% CO_2_) for 48 hours. After incubation, the area of gels was measured using ImageJ. The area of gels without a mixture of cells was set at 100%. The potential miRNA targets in the 3’ UTRs of *Col1a1, Col3a1* and *Tnxb* were predicted by TargetScan (http://www.targetscan.org/vert_80/) and miRWalk (http://mirwalk.umm.uni-heidelberg.de/) programs. The 3’ UTRs of *Col1a1, Col3a1* and *Tnxb* were PCR amplified using mouse genomic DNA and cloned into the pMIR-REPORT-LUC reporter vector (Ambion). The primers for PCR amplification include: forward 5 ATCGACTAGTTTTGGAGCCAGGCAGGGTCAC and reverse 5 AGCTAAGCTTTGGTCTAGGGAGCATCTCAGC for *Col1a1*, forward 5 ATCGACTAGTCACCCAATACAGGTCAAATGC and reverse 5 AGCTAAGCTTTATGGCTTGAAT GAAGGTACC for *Col3a1*, and forward 5 ATCGACTAGTTGCGACCCAGAAACTTCCAGG and reverse 5 AGCTAAGCTTCAGTTTCTCCTTTATTGCTCC for *Tnxb*. MiRNA mimics were purchased from Sigma (miR-98, #HMI0982; miR-30c, #HMI0458; miR-29a, #HMI0434; miR-196b, #HMI0325; miR-148a, #HMI0237; miRNA negative control, #HMC0002). The reporter constructs with different miRNA mimics (including a negative control miRNA) were co-transfected into tsA58-AVM cells using lipofectamine2000 (Invitrogen) as described previously (Peng et al., 2014). To normalize the transfection efficiency, a plasmid expressing a lacZ reporter driven by a constant promoter was included in the co-transfection. Luciferase activities were measured using the Luciferase Assay System (Promega, #E1500) and the Agilent Bio Tek Synergy 2 multi-mode microplate reader. β-galactosidase assays were performed with the same cell extract using the β-Galactosidase Enzyme Assay System with Reporter Lysis Buffer (Promega, #E2000). The luciferase activity was normalized against the lacZ activity of each culture. To examine expression of miRNAs through qRT-PCR, miRNA was purified from GFP+ cells of mitral leaflets or tricuspid leaflets from E18.5 *Nfatc1-Cre;mTmG* embryos using mirVana miRNA isolation kit (Ambion, #AM1560). Leaflets were digested with .25% trpsin, and GFP+ cells (endocardial + mesenchymal cells) were isolated using a FACS Aria II sorter (BD BioSciences, performed by the UAB Comprehensive Flow Cytometry Core facility). A total of 4 samples for each genotype were analyzed. For each sample, 10-15 leaflets were pooled together for RNA extraction. CDNA synthesis and qRT-PCR were performed using miRCURY LNA miRNA PCR Starter kit (Qiagen, #339320) according to the manufacturer’s instructions. MiRNA-specific PCR primers were purchased from Qiagen (miR-98-5p, #YP00204640; Let-7a-5p, #YP00205727; Let-7b-5p, #YP00204750; Let-7c-5p, #YP00204767; miR-29a-3p, #YP00204698; miR-29b-3p, #YP00204679; miR-29c-3p, #YP00204729; miR-30a-5p, #YP00205695; miR-30b-5p, #YP00204765; miR-30c-5p, #YP00204783). The level of miRNA expression was normalized against Snord65 (Qiagen, #YP00203910) expression.

### 6. ScRNA-Seq and trajectory analyses

Mitral leaflets were dissected from E17.5 control and mutant embryos and digested with 0.05% Trypsin to obtain single cells. For each genotype, samples were pooled from 3-4 hearts. To collect viable single cells, 7-amino-actinomycin (7-AAD, ThermoFisher, for labeling nonviable cells) stained cells were sorted using a FACS Aria II sorter (BD BioSciences). Preparation of the single cell transcriptome libraries was performed using 10xGenomics Single Cell 3□ v3 Reagent Kits, following the manufacturer’s instructions. The prepared libraries were sequenced using Illumina NextSeq500 with the expectation of at least 2,000 total reads per cell. The 10X Genomics Cellranger software (version 6.0.2), mkfastq, was to create the fastq files from the sequencer. After fastq file generation, Cellranger count was used to align the raw sequence reads to the mouse reference genome. The matrix tables created by the count was then loaded into the R package Seurat (version 4.0.5) which allows for selection and filtration of cells based on quality control metrics, data normalization and scaling, and detection of highly variable genes (Butler et al., 2018). We followed the Seurat vignette (https://satijalab.org/seurat/articles/pbmc3k_tutorial.html) to create the Seurat data matrix object. In brief, we kept all genes expressed in more than three cells and cells with at least 200 detected genes. Cells with mitochondrial gene percentages >10% and unique gene counts > 5000 or < 200 were discarded. The data were normalized using Seurat’s NormalizeData function, which uses a global-scaling normalization method, LogNormalize, to normalize the gene expression measurements for each cell to the total gene expression. The result is multiplied by a scale factor of 1e4 and the result is log-transformed. Highly variable genes were then identified using the function FindVariableFeatures in Seurat. We also regressed out the variation arising from library size and percentage of mitochondrial genes using the Seurat function ScaleData. We performed principal component analysis (PCA) of the variable genes as input and determined significant principal components on the basis of the Seurat JackStraw function. The first 50 principal components were selected as input for UMAP using FindNeighbors, FindClusters (resolution = 0.8) and RunUMAP in Seurat. To identify differentially expressed genes (DEGs) in each cell cluster, we used Seurat’s FindMarkers function on the normalized gene expression. To identify single-cell trajectory, the Seurat object was converted to a Monocle3 (Trapnell et al., 2014) object using the ‘as.cell_data_set’ function in Seurat. The cells were clustered to determine partitions using Monocle3 ‘cluster_cells’ function. The next step was to learn the trajectory graph using ‘learn_graph’. The cells were then ordered in pseudotime using ‘order_cells’.

### 7. Overall statistical analyses

All data are shown as mean ± SEM. Unpaired Student’s *t*-test was used to compare between control and mutant groups with *p*<0.05 considered statistically significant. P values for each dataset are provided in the corresponding figures or figure legends. For detection of differentially expressed genes between control and mutant cells in scRNA-Seq analysis, the Benjamini-Hochberg adjusted P values were calculated.

## Acknowledgments

We thank Dr. B. Zhou (Elbert Einstein Medical School) for providing *Nfatc1-Cre* mice. We thank the UAB Genomics Core Facility for performing deep-Seq and Sanger Sequencing. We thank the members of Dr. Jiao’s lab for their suggestions that helped support this project.

## Supplementary figures

**Sup. Figure 1.**
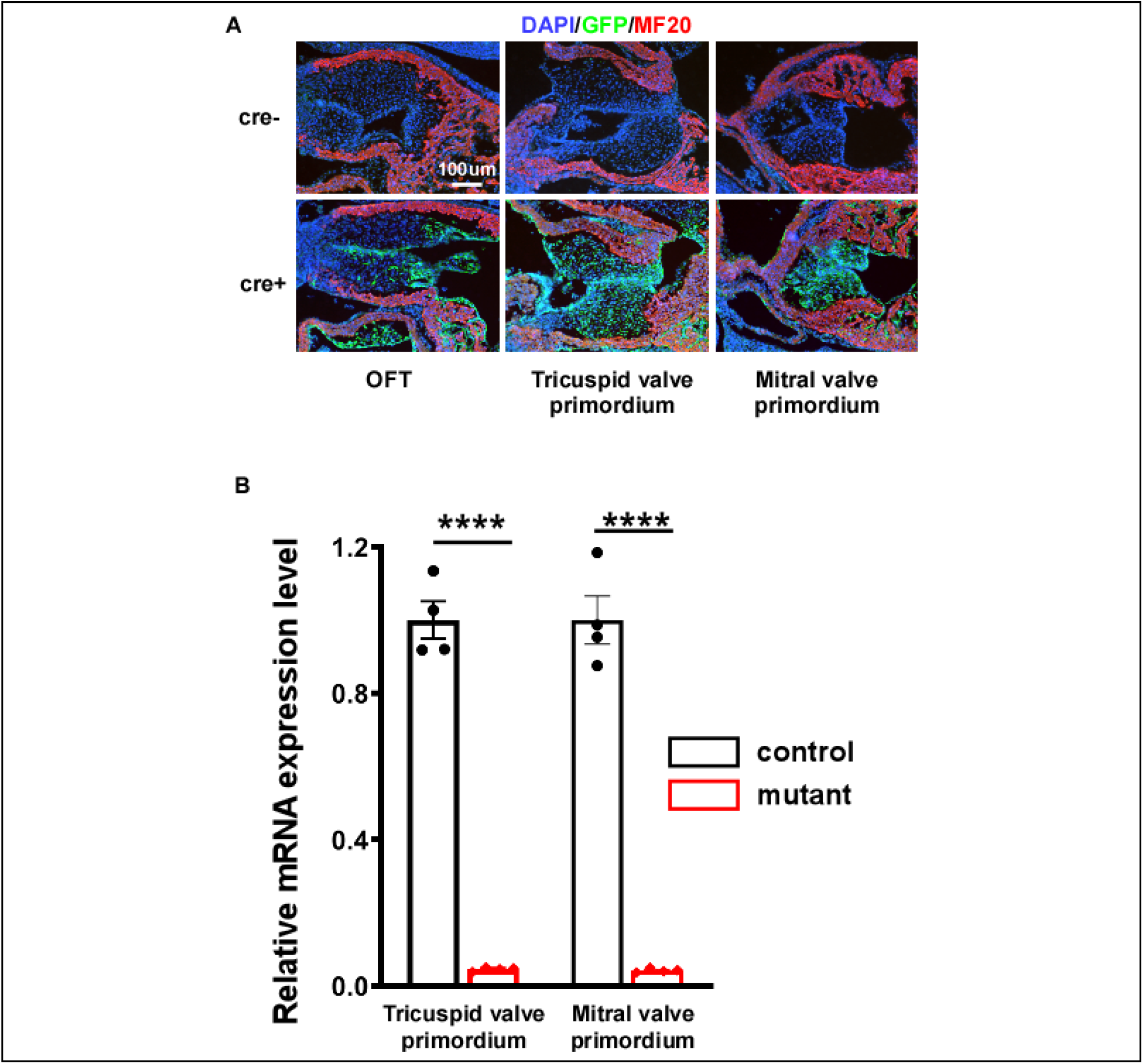
**(A)** *Nfatc1-Cre* heterozygous male mice were crossed with mTmG/mTmG reporter mice to obtain Cre+ and Cre-negative embryos at E11.5. These embryos were then sagittally sectioned and stained with antibodies against cardiac myosin heavy chain (MF20, red) and GFP (green). In Cre+ embryos, GFP signals (indicative of Cre activity) were readily detected in the proximal region of outflow tract (OFT) cushions, tricuspid valve primordium and mitral valve primordium. **(B)** Cells were isolated tricuspid valve primordium and mitral valve primordium of control (*Nfatc1-Cre;Dicer1*^*+/+*^*;mTmG*) and mutant (*Nfatc1-Cre;Dicer1*^*loxp/loxp*^*;mTmG*) embryos at E11.5) isolated through laser capturing (Leica LMD6, UAB Laser Microdissection Facility). Total RNA were isolated from these cells followed by qRT-PCR using primers corresponding the exon 3 (Peng et al., 2014), which would be removed upon CRE-mediated recombination. The control level was set at 1.0. Data were averaged from 4 independent samples and were shown as mean±SEM. ****: *p<0*.*0001*, unpaired, two tailed Student’s t test.

**Sup. Figure 2.**
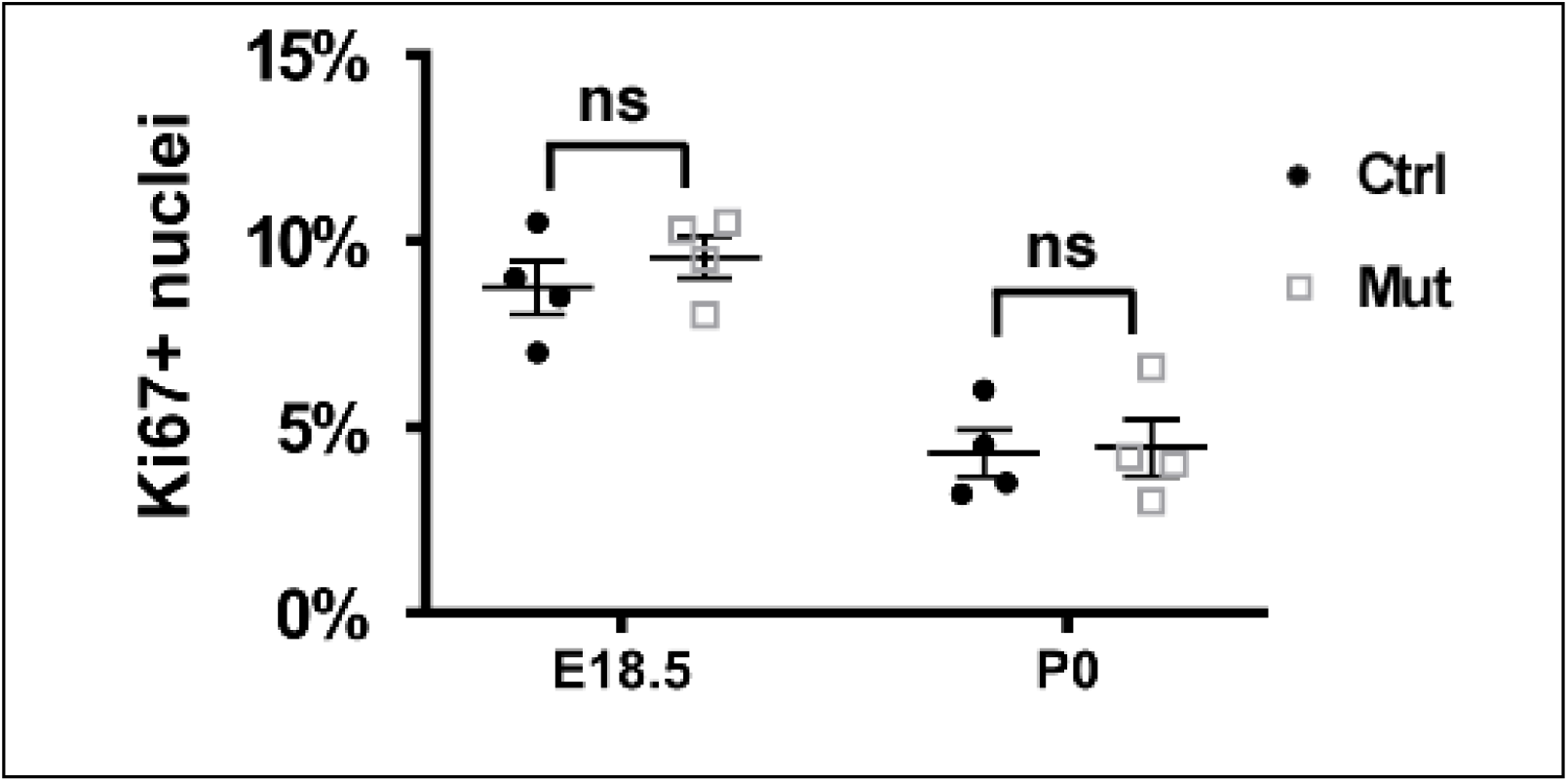
Quantification of cell proliferation in mitral valves at E18.5 and P0. Cardiac sections were stained with antibodies against Ki67 and myosin heavy chain (mf20) as shown in Figure 2C. The percentage of Ki67+ nuclei among total nuclei was determined from 4 independent hearts. For each heart, at least 3 sections were included and for each section at least 100 nuclei were counted. Ns: no significant difference, unpaired, two tailed Student’s t test.

**Sup. Figure 3.**
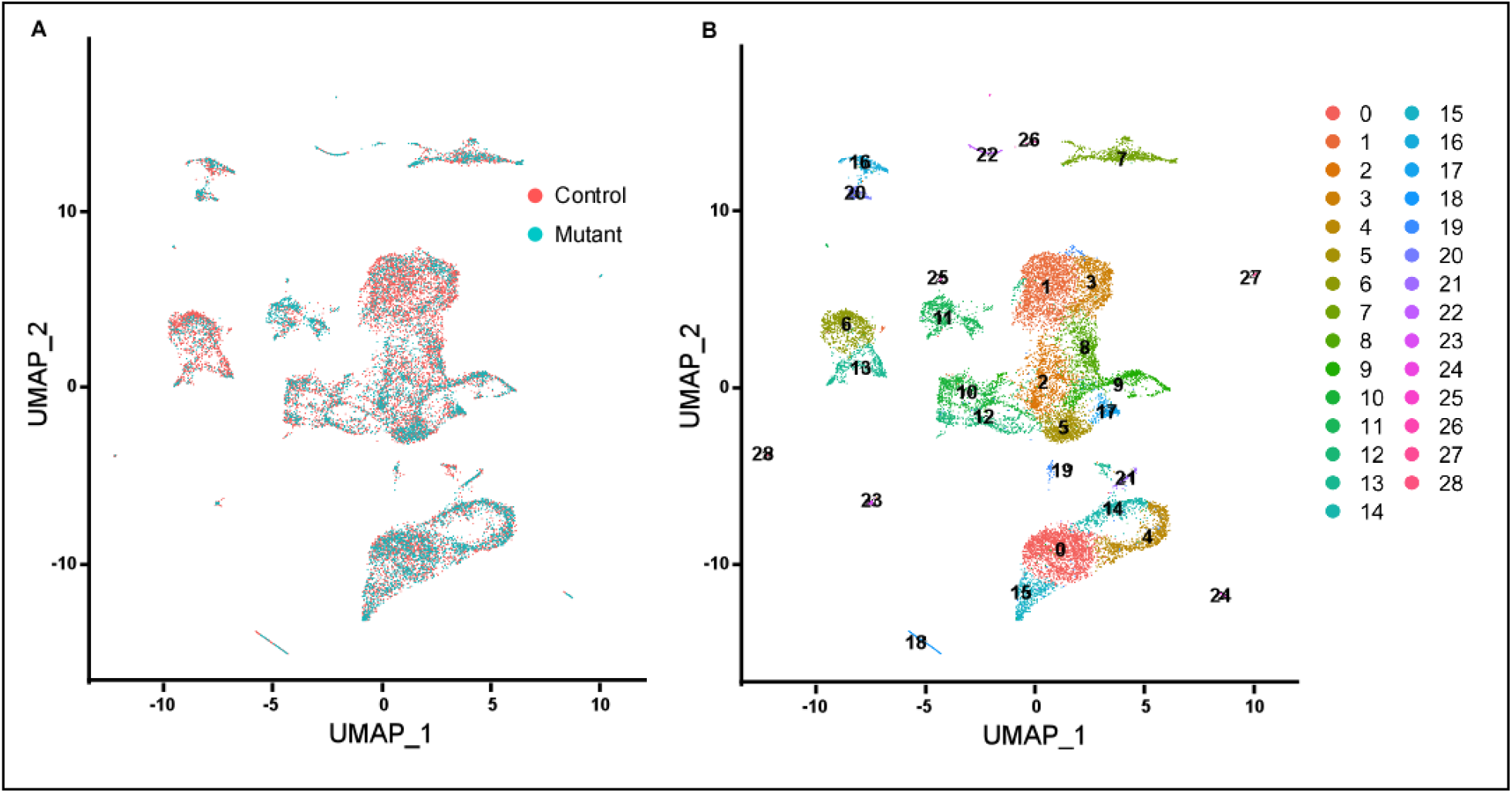
**(A)** The UMAP chart shows the distribution of control (red) and mutant (blue) cells across all clusters. **(B)** The UMAP chart shows that the cells isolated from mitral valve regions can be grouped into 29 clusters.

**Sup. Figure 4.**
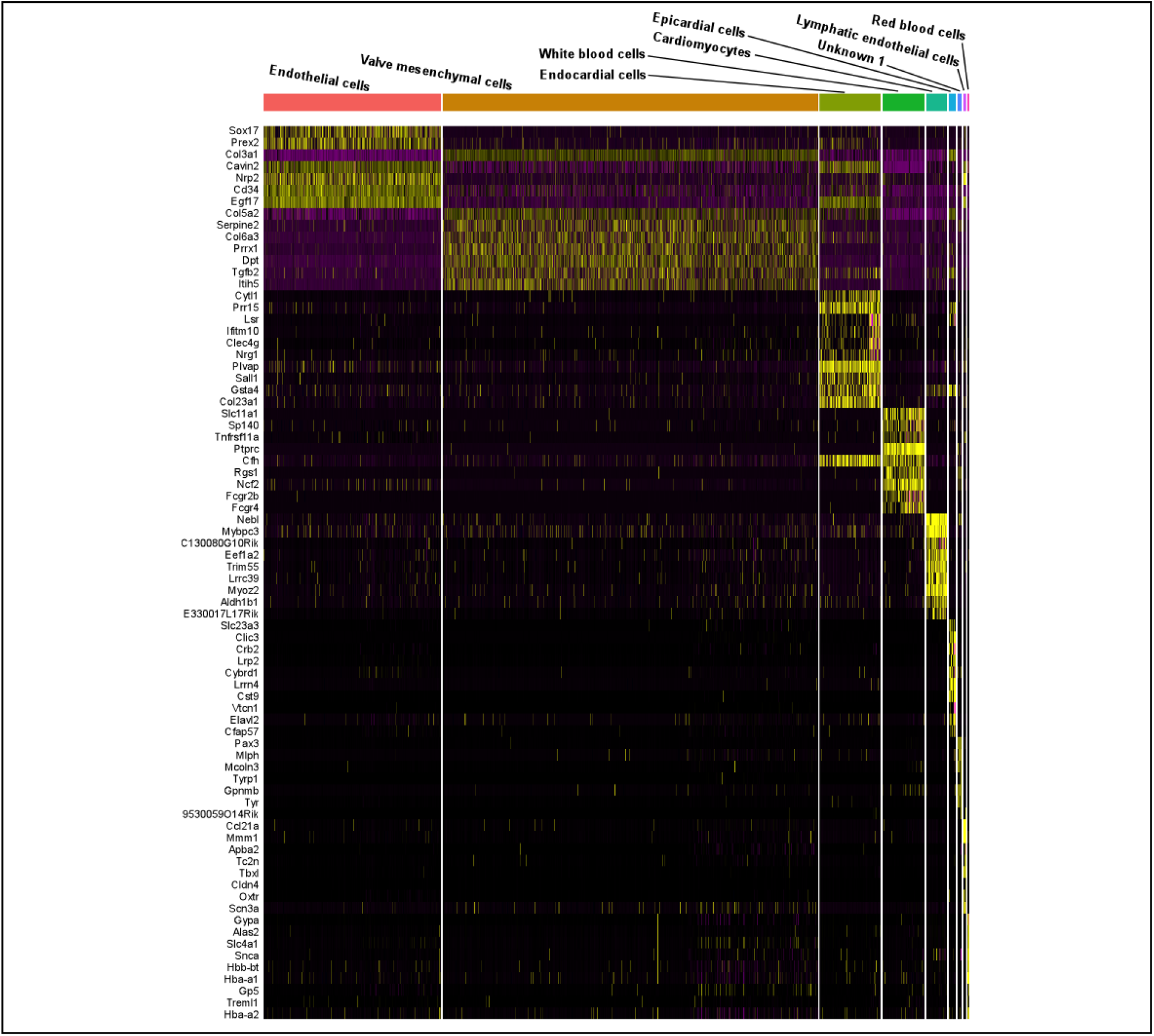
Expression-based heat map of the top 10 genes that are differentially expressed in each cluster of cells.

**Sup. Figure 5.**
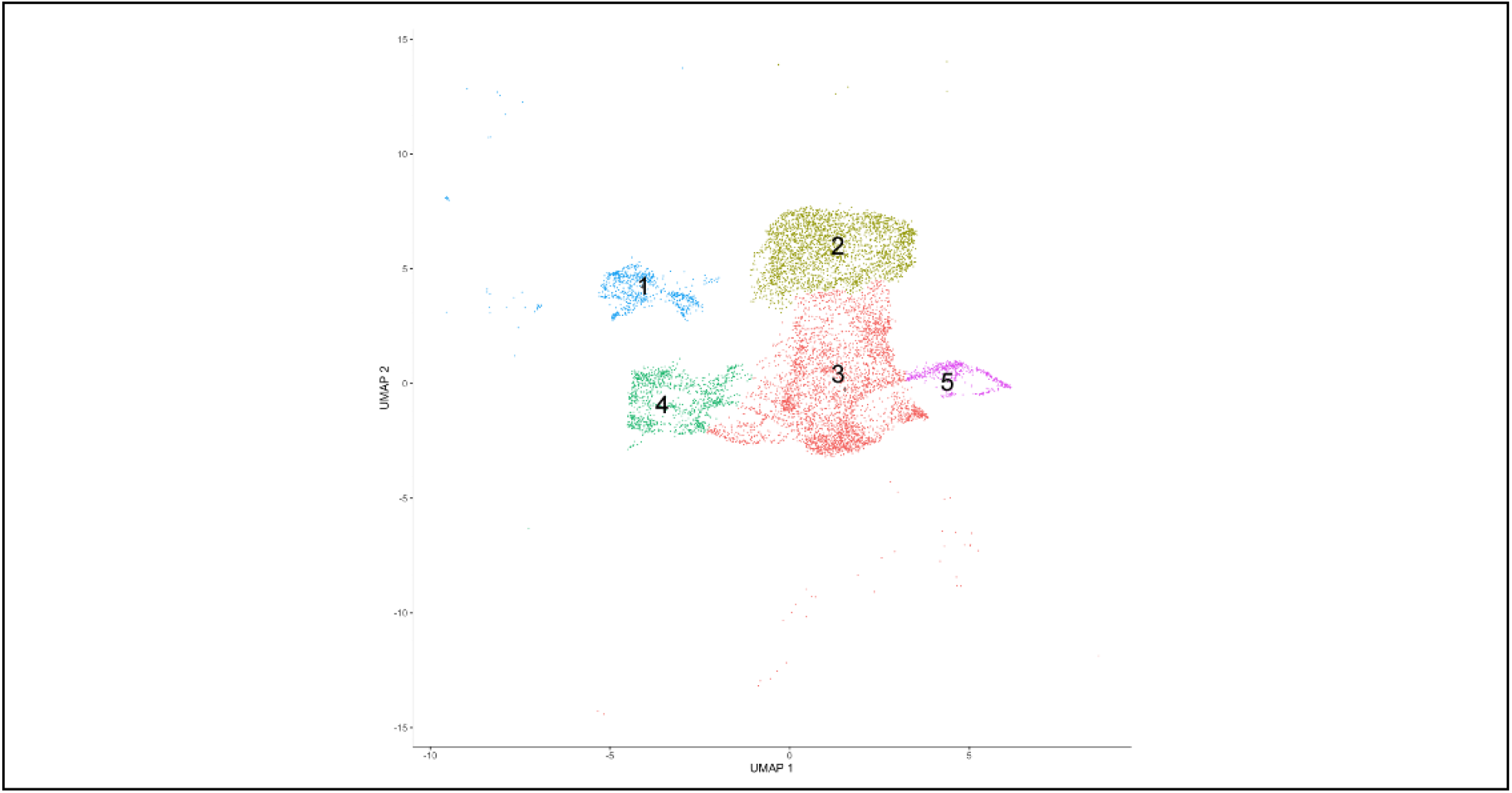
Cushion mesenchymal cells can be further grouped into 5 subclusters using Monocle 3.0.

**Sup. Figure 6.**
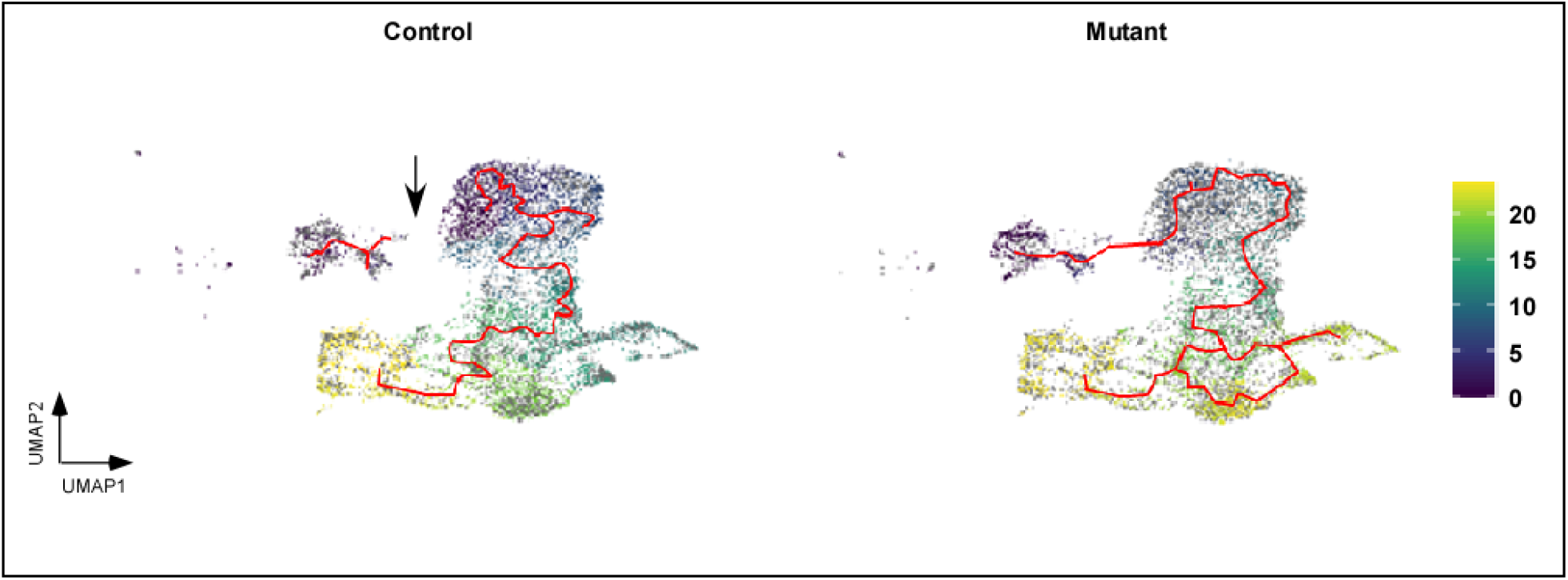
Pseudotime analysis of control and mutant valve mesenchymal cells separately. The time order is shown with colors. The red line indicates the trajectory pathways. The pathway between subcluster 1 and other subclusters is broken as indicated by the arrow.

**Sup. Figure 7.**
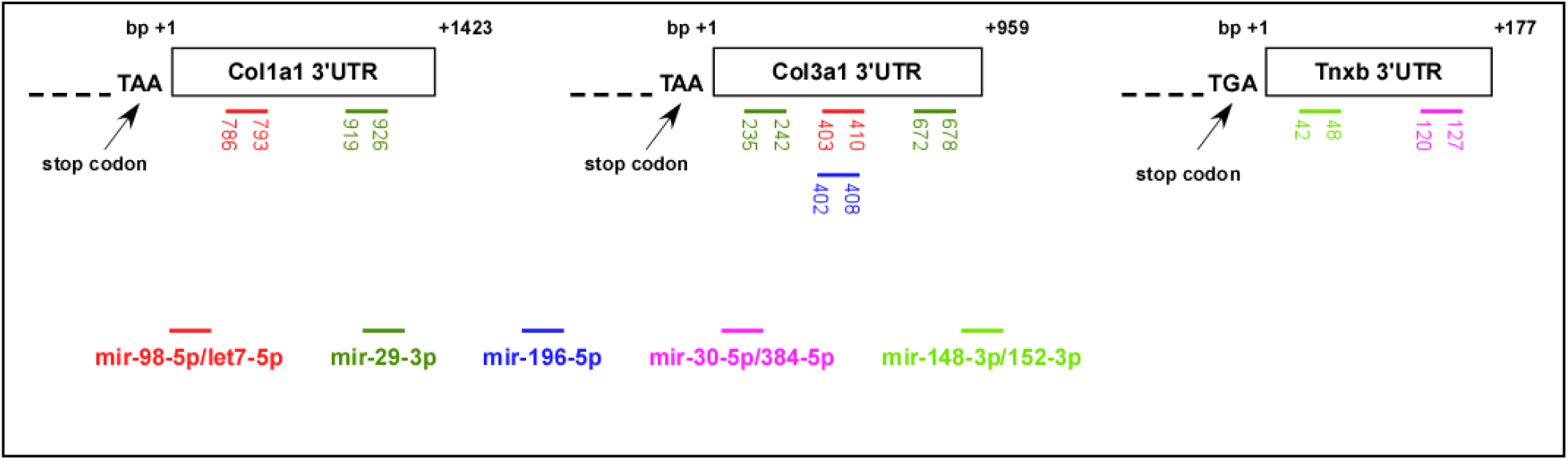
The 3’ UTR regions of *Col1a1, Col3a1* and *Tnxb* contain potential target sequences of multiple miRNAs as predicted from - TargetScan and miR Walk. Only the sequences that are predicted by both programs and are highly conserved among mammals are shown.

**Sup. Figure 8.**
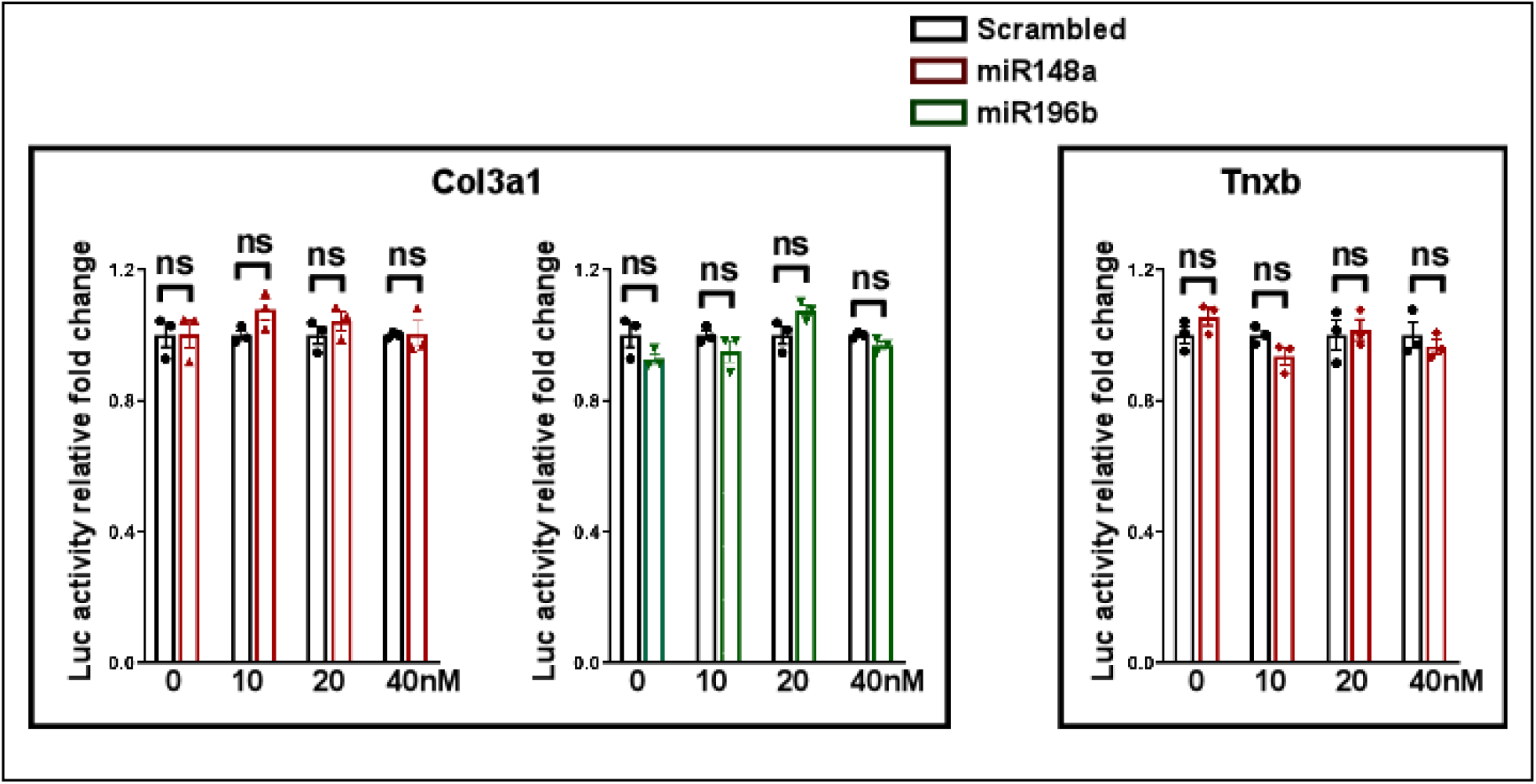
MiR148a and miR196 mimics failed to alter the activity of reporters generated using 3’ UTR of *Col3a1* and *Tnxb*. Data are averaged from 3 independent cultures and are shown as mean ± SEM. Ns: no significant difference, unpaired, two tailed Student’s t test.

